# Preclinical Synergistic Combination Therapy of Lurbinectedin with Irinotecan and 5-Fluorouracil in Pancreatic Cancer

**DOI:** 10.1101/2023.09.13.557653

**Authors:** Tej Tummala, Ashley Sanchez Sevilla Uruchurtu, Arielle De La Cruz, Kelsey E. Huntington, Andrew George, Nicholas R. Liguori, Leiqing Zhang, Lanlan Zhou, Abbas E. Abbas, Christopher G. Azzoli, Wafik S. El-Deiry

## Abstract

Pancreatic cancer is a devastating disease with a poor prognosis. Novel chemotherapeutics in pancreatic cancer have shown limited success, illustrating the urgent need for new treatments. Lurbinectedin (PM01183; LY-01017) received FDA approval in 2020 for metastatic small cell lung cancer on or after platinum-based chemotherapy and is currently undergoing clinical trials in a variety of tumor types. Lurbinectedin stalls and degrades RNA Polymerase II and introduces breaks in DNA, causing subsequent apoptosis. We now demonstrate lurbinectedin’s highly efficient killing of human-derived pancreatic tumor cell lines PANC-1, BxPC-3, and HPAF-II as a single agent. We further demonstrate that a combination of lurbinectedin and irinotecan, a topoisomerase I inhibitor with FDA approval for advanced pancreatic cancer, results in synergistic killing of pancreatic tumor cells. Western blot analysis of combination therapy indicates an upregulation of γH2AX, a DNA damage marker, and the Chk1/ATR pathway, involved in replicative stress and DNA damage response. We further demonstrate that the triple combination between lurbinectedin, irinotecan, and 5-fluorouracil (5-FU) results in highly efficient killing of tumor cells. Our results are developing insights regarding molecular mechanisms underlying therapeutic efficacy of a novel combination drug treatment for pancreatic cancer.

## Introduction

Pancreatic cancer is a devastating disease with a poor prognosis and has an overall 5-year relative survival rate of 12% [1]. Given current screening methods, patients are often diagnosed with metastasized stage IV pancreatic cancer with no option for surgery [2]. Less than 20% of pancreatic tumors are resectable, and patients with successful resection have a 5-year survival rate of approximately 20% [3]. A combination therapy used in pancreatic cancer is FOLFIRINOX, a combination of folinic acid (leucovorin calcium), fluorouracil, irinotecan, and oxaliplatin. While somewhat difficult to tolerate, FOLFIRINOX has been shown to be highly effective in certain patient populations compared to gemcitabine monotherapy [4]. The FOLFIRINOX regimen has comparable efficacy to gemcitabine plus abraxane in advanced pancreatic cancer [5]. Despite developments in novel chemotherapeutics and targets for pancreatic cancer, current therapies have showed limited success. Thus, discovering new chemotherapeutics and combination therapies in pancreatic cancer is essential.

Lurbinectedin (ZEPZELCA™), developed by PharmaMar yet frequently distributed and advertised by the U.S. based company Jazz Pharmaceuticals, received Food and Drug Administration (FDA) approval in the United States in June 2020 for adult patients with metastatic small-cell lung cancer with progression of the disease on or after platinum-based chemotherapy [6]. Lurbinectedin is a novel tetrahydroisoquinoline shown to degrade RNA-Polymerase II and covalently bind to guanine-rich sequences in the minor groove of DNA [7]. Following binding to the minor groove, lurbinectedin forms adducts which can induce double-strand breaks, leading to subsequent apoptosis [8]. It can further preferentially bind to DNA at transcription factor recognition sites, and it is hypothesized this could prevent transcription factors from appropriately binding to chromatin [9]. While the mechanism of lurbinectedin is still being explored in the tumor and tumor microenvironment, preclinical models indicate that lurbinectedin reduces tumor-associated macrophages (TAMs) and reduces the production of certain inflammation and growth factors, including CCL2 and VEGF [10].

Irinotecan (IRT), introduced to the clinic in 1998, is a cytotoxic prodrug for SN38, which plays a role in the inhibition topoisomerase I. SN38 stabilizes the complex formed by topoisomerase-I and DNA, inducing single-strand breaks. Eventually, these single-strand breaks yield double strand breaks due to replication fork collapse, triggering subsequent apoptosis [11]. IRT has been used to treat a variety of solid tumors, including but not limited to lung cancer, colorectal cancer, and pancreatic cancer [12]. It is frequently used in combination therapies such as FOLFIRI and FOLFIRINOX or FOLFOXIRI for many metastasized or advanced gastrointestinal (GI) solid tumors [13]. Given it was approved over twenty-five years ago and still plays a major role in many standard chemotherapeutic regimens, irinotecan has established itself as an essential GI anticancer drug [14].

5-Fluorouracil (5-FU), first discovered in 1957, is used in a variety of tumor types and has been shown to have many potential mechanisms of antitumor activity [15, 16]. 5-FU is capable of interfering with the biosynthesis of nucleic acids, and it can be misplaced in RNA and DNA instead of uracil and thymine due to its structural similarity to pyrimidines [17]. This substitution has been shown to induce cytoxicity and cell death. It is used in many combination therapies in GI cancers alongside irinotecan, including the FOLFOX, FOLFIRI, FOLFOXIRI and FOLFIRINOX regimens [18].

Interest in lurbinectedin is rising. The drug is being tested in both preclinical and clinical settings as a single agent or in combination with other chemotherapeutics in a variety of tumor types. There is currently an ongoing phase II trial of lurbinectedin monotherapy in a variety of patients with advanced solid tumors, including advanced pancreatic cancer [19]. There is also an ongoing phase I/II clinical trial combining lurbinectedin and irinotecan in a variety of tumor types, included pancreatic adenocarcinoma, that is estimated to be completed in late 2023 [20]. Further, there is an ongoing phase III clinical trial combining lurbinectedin and irinotecan in patients with relapsed small-cell lung cancer [21], indicating the combination has potential in other tumor types. However, there is no current ongoing clinical trial with lurbinectedin monotherapy or combination therapy with irinotecan specifically in pancreatic adenocarcinoma.

Our group further explored the combination of lurbinectedin and irinotecan in a preclinical setting using pancreatic cancer cell lines. We report synergistic killing of such cells and changes in protein and cytokine expression under single agent and combination therapies. We report highly efficient killing of pancreatic cancer cells including with the triple combination between lurbinectedin, irinotecan, and 5-FU.

## Materials and Methods

### Cell Lines and Culture Conditions

PANC-1, HPAF-II, and BxPC-3 are pancreatic cancer cell lines obtained from American Type Culture Collection (ATCC). PANC-1 was isolated from the pancreatic duct of a 56-year-old white male with epithelioid carcinoma. HPAF-II was isolated from a 44-year-old white male patient with metastasized pancreatic adenocarcinoma. BxPC-3 was isolated from the pancreas tissue of a 61-year-old female patient with adenocarcinoma. While all three cell lines have different genetic profiles, KRAS was activated in PANC-1 and HPAF-II, whereas BxPC-3 was wild-type. *TP53* mutations are found in all cell lines [22]. All cell lines were cultured in Dulbecco’s Modified Eagle’s Medium (DMEM) media with an additional 10% FBS and 1% penicillin-streptomycin solution added. Cells were all incubated at 37°C and monitored consistently.

### Therapeutic agents

Lurbinectedin was obtained through MedChem Express (Catalog #HY-16293). Irinotecan was obtained through Sigma (Catalog #1406-50MG). 5-FU was obtained through Selleck Chemicals (Catalog #S1209). In accordance with the manufacturer’s recommendations, each of the above therapeutics were stored at -20°C.

### Cell Viability Analysis and Combination Synergy Score Analysis

Cell viability was determined via a CellTiter-Glo (CTG) assay. Approximately 3,000 cells were plated per well in a black 96-well plate and left to incubate overnight. After 12-24 hours allowing for cells to adhere, cells were treated with lurbinectedin, irinotecan, or 5-FU at increasing concentrations to determine single drug IC-50s. Drug concentrations ranged from a maximum value to a control, diluted via 1:2 serial dilution. The well plates were re-incubated for 72 hours. After the re-incubation period, CTG technology indicated cell viability by luminescent detection of ATP. The percentage of live cells was calculated and analyzed relative to the control. IC-50 values at 72 hours were determined for the three drugs in the three cell lines (PANC-1, BxPC-3, and HPAF-II).

Similarly, combination drug treatments of both two and three drugs were performed via a CellTiter-Glo assay. Approximately 3,000 cells were plated per well in a 96-well plate and left to incubate overnight. All three previously mentioned cell lines were used. The following day, cells were treated with lurbinectedin + irinotecan or lurbinectedin + irinotecan + 5-FU at increasing drug concentrations such that the leftmost upper corner well was a control in every plate. CTG was used to evaluate cell viability. Combination index analysis was performed via the combination index software SynergyFinder, which calculated synergy scores according to an HSA reference model [23]. HSA calculates synergy scores based on the percent inhibition above the maximum single drug response at a given concentration. The website of the manufacturer indicates a synergy score great than 10 is synergistic killing of cells whereas a synergy-score lower than -10 indicates antagonistic killing of cells. From -10 to 10, there is likely an additive interaction between the drugs. Synergistic killing refers to a greater than additive interaction between the drugs.

### Western Blot Analysis

Western blot analysis was performed to analyze the effects of lurbinectedin, irinotecan, and combination in PANC-1, BxPC-3, and HPAF-II pancreatic cancer cell lines. Western blot analysis demonstrated changing protein expression levels, indicating the molecular mechanisms underlying synergy. Cells were plated at approximately 500,000 cells per well in a 6-well plate. After 12-24 hours, giving sufficient time for cells to adhere, the cells were treated with control, single, or combination therapies. After 48 hours of incubation, cells were harvested from the well-plates using a scraper. Cells were lysed with RIPA buffer treated with a protease and phosphatase inhibitor. Proteins were quantified via a BCA assay using the Pierce BCA Protein Assay Kit obtained from Life Technologies (Catalog #23225). Cell lysates were run through a 4-12% SDS-PAGE gel electrophoresis at 140-170V, and subsequently transferred to a PVDF membrane. The PVDF membrane was blocked for 30 minutes with 5% non-fat milk (milk powder diluted in TBST). Primary antibodies were diluted at their respective manufacturer’s suggested Western Blot dilution in 5% non-fat milk. Following blocking, membranes were incubated in primary antibodies overnight at 4°C. Invitrogen Goat anti-Rabbit IgG (HRP #31460) and Goat anti-Mouse IgG (HRP #31430) were used as secondary antibodies and diluted 1:5000 in 5% nonfat milk. Each membrane was probed using ECL with an appropriate loading control, which was either vinculin or actin.

### Cytokine Profiling Analysis

HPAF-II and BxPC-3 cell lines were plated at a density of 75,000 cells/well in a 24 well plate in 0.5 mL of DMEM media. After 16 hours, allowing sufficient time for cells to adhere, cells were treated with no drug, lurbinectedin, irinotecan, or combination, and re-incubated for 48 hours. Following this incubation, media was harvested, and cytokine levels were analyzed using a Luminex Cytokine Detection Assay Panel.

## Results

### Lurbinectedin is a highly potent cytotoxic agent in pancreatic adenocarcinoma cell lines

Given the documented low IC-50s of lurbinectedin in small cell lung cancer across the literature, it was hypothesized that lurbinectedin may be an efficient killer of pancreatic tumor cells as well. To examine this, HPAF-II, BxPC-3, and PANC-1 human derived pancreatic adenocarcinoma cells were cultured (**Figure 1**).

**Figure 1.**
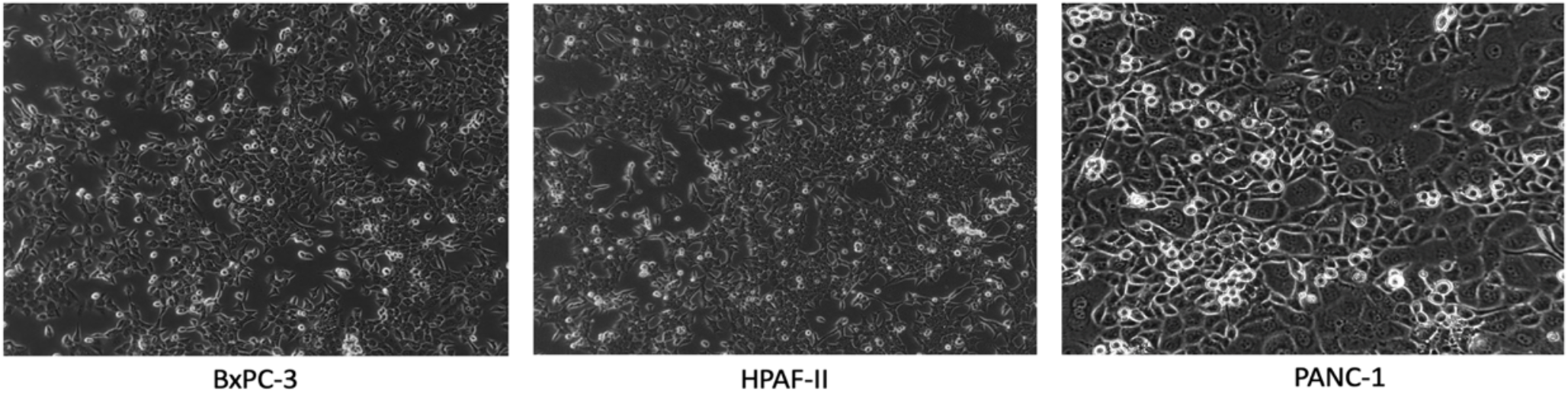
Microscopic images of human derived pancreatic cancer cell lines. The above images are microscopic images of the pancreatic cell lines BxPC-3, HPAF-II, and PANC-1. The cell lines exhibit the expected morphology seen on the ATCC website from which they were obtained.

After plating three human-derived pancreatic cancer cell lines **Figure 1**, cells were treated with increasing concentrations of lurbinectedin as a single agent (**Figure 2**). Further, cells were treated with irinotecan and 5-FU as single agents, as these drugs are commonly used in modern chemotherapeutic regimens for patients with pancreatic adenocarcinoma (**Figure 2**).

**Figure 2.**
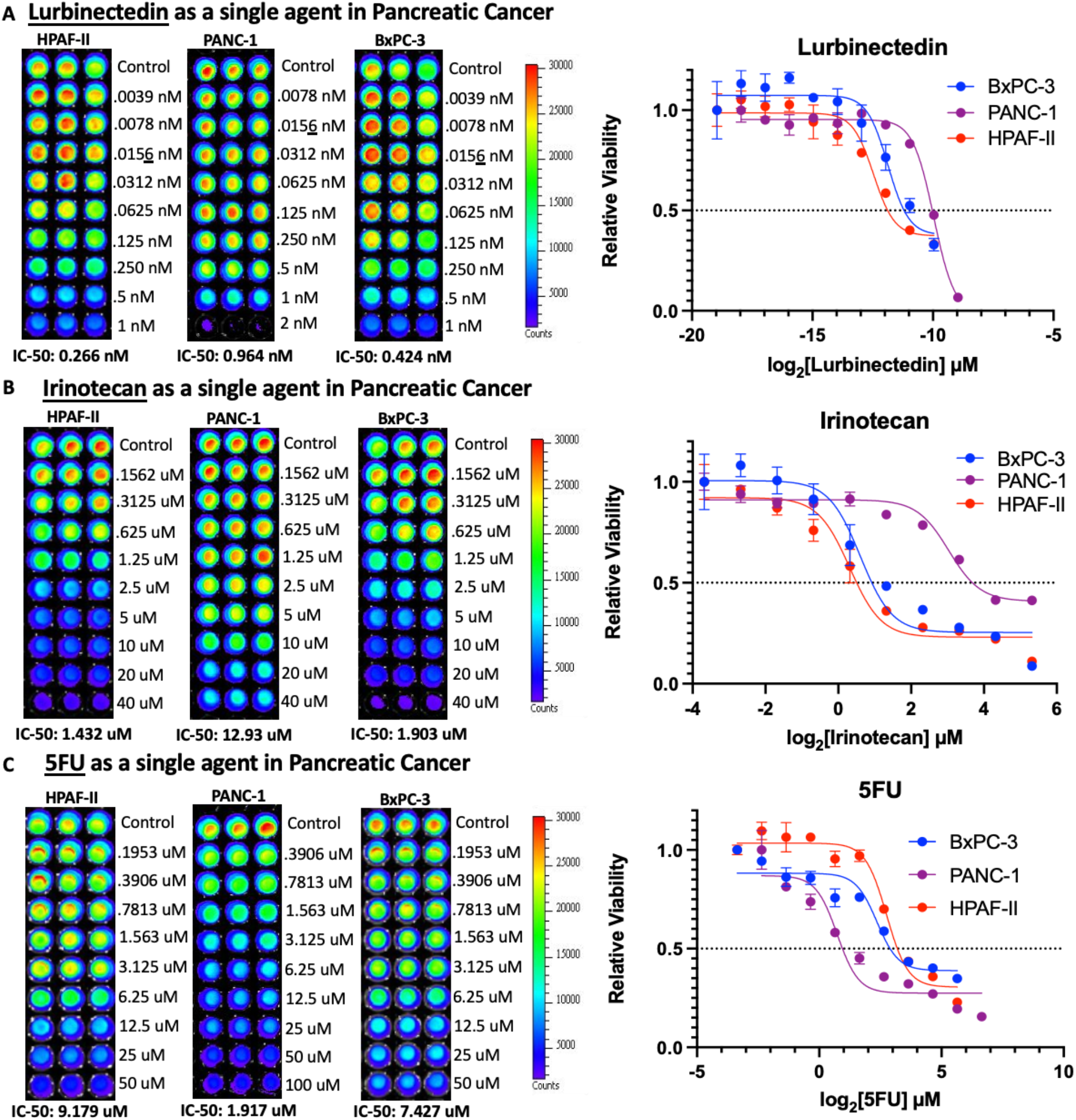
Lurbinectedin, irinotecan, and 5-FU as single agents in pancreatic cancer. CellTiter-Glo technology utilizes luminescent detection of ATP, and thus brighter luminescence corresponds to higher cell viability. (**a**) Lurbinectedin exhibits sub-nanomolar 72-hour IC-50s in HPAF-II, PANC-1, and BxPC-3 cell lines, demonstrating highly efficient killing of pancreatic cancer cells *in vitro*. The most sensitive cell line was HPAF-II, with an IC-50 value of 0.266 nM, whereases the most resistant cell line was PANC-1 with an IC-50 of 0.964 nM. (**b**) Irinotecan demonstrates 72-hour IC-50s mostly in the low micromolar range. HPAF-II is the most sensitive cell line whereas PANC-1 was the most resistant cell line, with IC-50s corresponding to 1.432 μM and 12.93 μM, respectively. (**c**) 5-Fluoruacil also demonstrates 72-hour IC-50s in the relatively low micromolar range. PANC-1 is the most sensitive cell line and HPAF-II was the most resistant cell line, corresponding to 1.917 μM and 9.179 μM, respectively.

### Lurbinectedin and irinotecan synergize in killing pancreatic adenocarcinoma cell lines

Given that lurbinectedin and irinotecan are currently in a variety of ongoing clinical trials in many tumor types, it was hypothesized that lurbinectedin and irinotecan may synergize in killing pancreatic adenocarcinoma cells, as irinotecan is frequently used in the clinic in a variety of combination therapies to treat pancreatic cancer.

Across all cell lines, there is a consistent trend of enhanced percent inhibition as each drug concentration increases (**Figure 3**). At sub-nanomolar concentrations of lurbinectedin and low micromolar concentrations of irinotecan, synergy scores greater than 15 are seen across all cell lines, indicating the potency of the combination in this tumor type. Once relatively strong synergism was established via viability assays, western blot analysis was performed to better understand the molecular mechanism underlying synergism.

**Figure 3.**
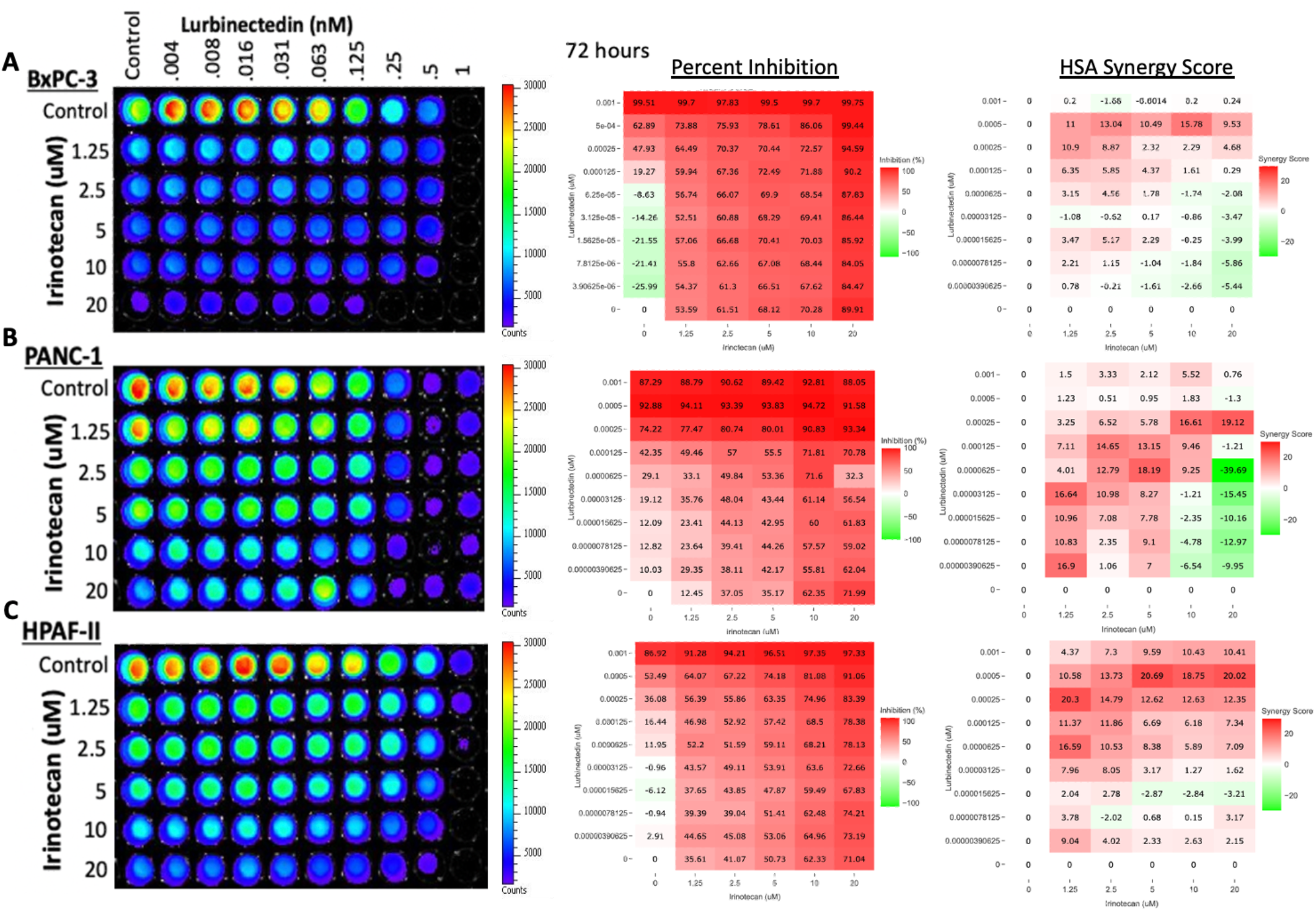
Combination therapy of lurbinectedin and irinotecan in BxPC-3, PANC-1, and HPAF-II results in efficient synergistic killing of pancreatic cancer cells. Lurbinectedin concentration ranges from 0-1 nM whereas irinotecan ranges from 0-20 μM. Percent inhibition is calculated relative to the control well (top left) of each plate. HSA synergy scores greater than 10 indicate synergistic killing of cells according to the website manufacturer. (**a**) In BxPC-3, the strongest synergism occurs at 0.5 nM of lurbinectedin and 10 μM of with an HSA synergy score of 15.78. At these concentrations, lurbinectedin and irinotecan as individual agents yield 62.89% and 70.28% inhibition, respectively, yet combined yield 86.06% inhibition. (**b**) In PANC-1, the strongest synergism occurs at 0.25 nM of lurbinectedin and 20 μM of irinotecan with an HSA synergy score of 19.12. At these concentrations, lurbinectedin and irinotecan individually yield 74.22% and 71.99% inhibition, respectfully, yet when combined, result in 93.34% inhibition. (**c**) In HPAF-II, the strongest synergism occurs at 0.5 nM of lurbinectedin and 5 μM of irinotecan, with an HAS score of 20.69. At these concentrations, lurbinectedin and irinotecan individually yield 53.49% and 50.73% inhibition, respectively, yet when combined result in 74.18% inhibition.

### DNA damage and replicative stress likely underlies synergism between lurbinectedin and irinotecan

Given that lurbinectedin is an RNA Polymerase II inhibitor shown to induce subsequent DNA damage and irinotecan is a topoisomerase I inhibitor which further causes DNA damage, it was hypothesized that the mechanism underlying synergism is likely rooted in genotoxicity. To test such a hypothesis, markers of DNA damage and other stress pathways were probed for (**Figure 4**).

**Figure 4.**
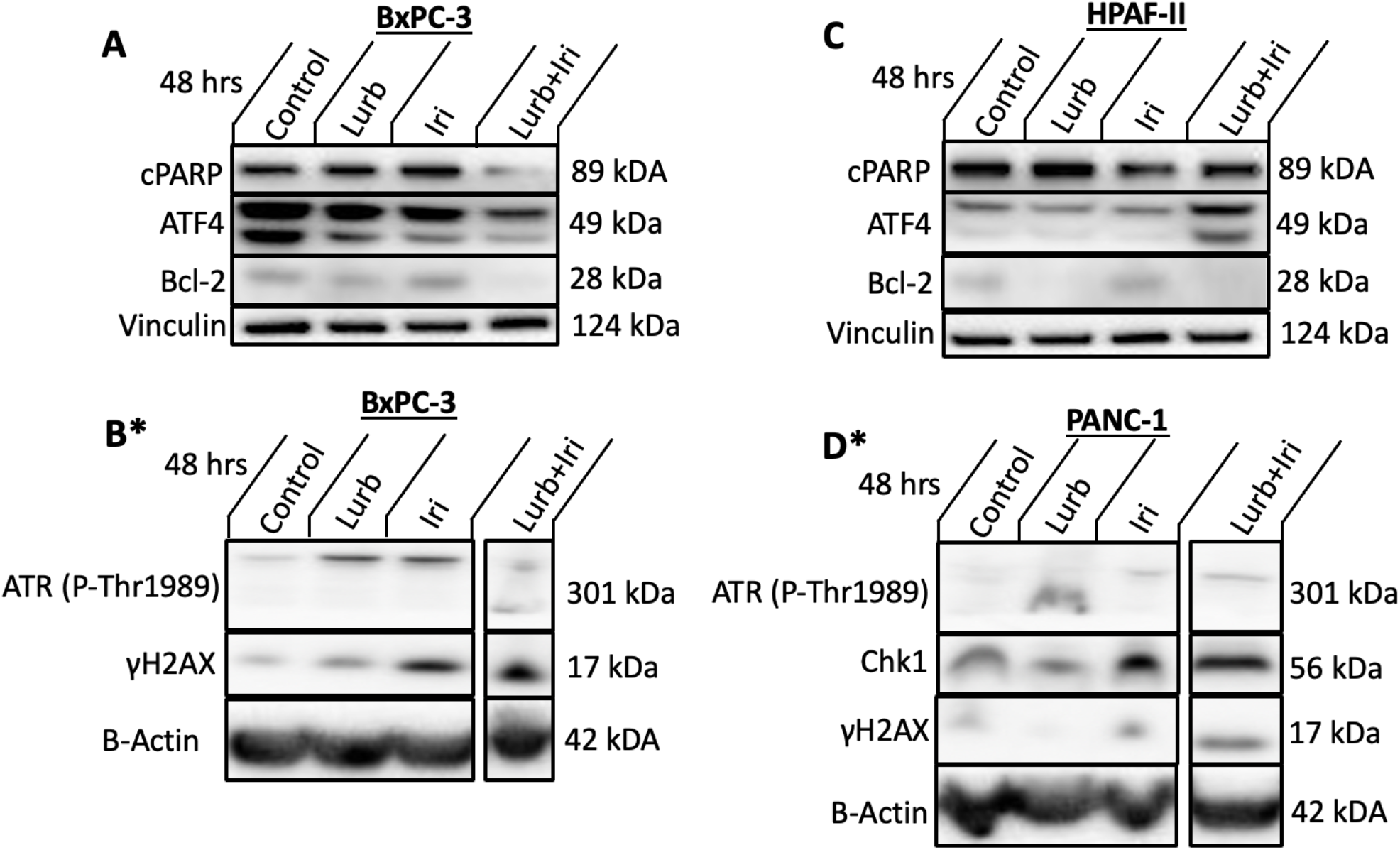
Western blot analysis of lurbinectedin and irinotecan as single agents and in combination in various pancreatic cancer cell lines. (**a**) & (**c**) cells were treated with either no drug, 0.5 nM of lurbinectedin, 10 μM of irinotecan, or the combination, for 48 hours. In (**b**) and (**d**), cells were treated with either no drug, 0.6 nM of lurbinectedin, 6 μM of irinotecan, or the combination. (**a**) & (**c**). In BxPC-3, there is a noticeable upregulation in γH2AX levels in both lurbinectedin and irinotecan-treated cells as single agents and in combination relative to the control group. There is an upregulation of ATR in the single agent treatment conditions and potentially a slight upregulation of ATR expression under combination conditions. Combination conditions also indicate a downregulation of Bcl-2. (**c**) HPAF-II, similarly, showed a downregulation of Bcl-2 in the combination conditions relative to the control. There is also a downregulation of ATF4. (**d**) In PANC-1, under single agent and combination conditions, there seems to be a slight upregulation of ATR. Chk1 expression is strongly upregulated in irinotecan and combination conditions. γH2AX are increased in combination conditions. *Some lanes were cropped out of the indicated gels due to irrelevance towards this paper, see Appendix A for the original western blot image.

Following western blot analysis, cytokine profiling analysis was performed to better understand the molecular mechanisms underlying synergism. A wide panel of cytokines were probed for, and fold change under different treatment conditions can be visualized in **Figure 5**.

**Figure 5.**
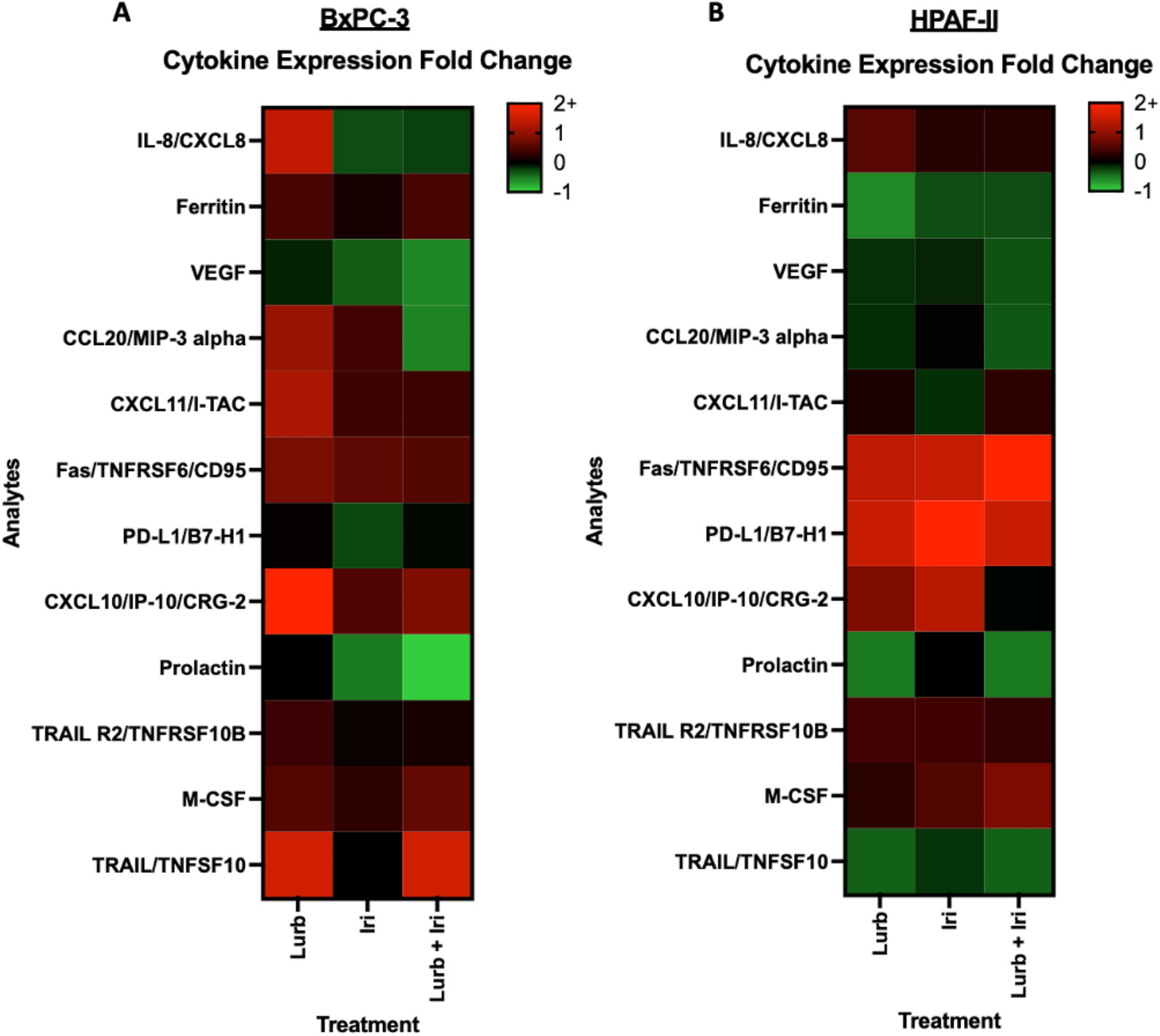
Cytokine profiling analysis shows upregulation and downregulation of a variety of cytokines under single agent and combination therapy conditions. **(a)** BxPC-3 cells and **(b)** HPAF-II cells were treated with either no drug, 0.25 nM of lurbinectedin, 2.5 μM of irinotecan, or the combination for 48 hours. Cytokine profiling analysis shows relatively large positive (upregulated: red) and negative (downregulated: green) fold-changes in a variety of cytokine markers relative to the control. Noticeably, monotherapy and combination therapy across both cell lines yielded a decrease in VEGF, which often plays a critical role in angiogenesis. Macrophage colony-stimulating factor (M-CSF) and Fas, a ligand involved in multiple apoptotic pathways, are upregulated in both monotherapy and combination therapy across both cell lines.

### Lurbinectedin, irinotecan, and 5-FU are a highly potent combination therapy in pancreatic adenocarcinoma cell lines

After establishing the synergism between lurbinectedin and irinotecan and investigating the molecular mechanisms underlying synergism, a triple combination of lurbinectedin, irinotecan, and 5-FU was examined to see if the presence of lurbinectedin could enhance the synergism between irinotecan and 5-FU, which is a drug combination present in many pancreatic cancer regimens (**Figure 6**). Given that lurbinectedin and irinotecan likely induce high levels of DNA damage, and 5-FU interferes with nucleic acid biosynthesis, it was hypothesized that the three drugs would act synergistically in killing pancreatic tumor cells.

**Figure 6.**
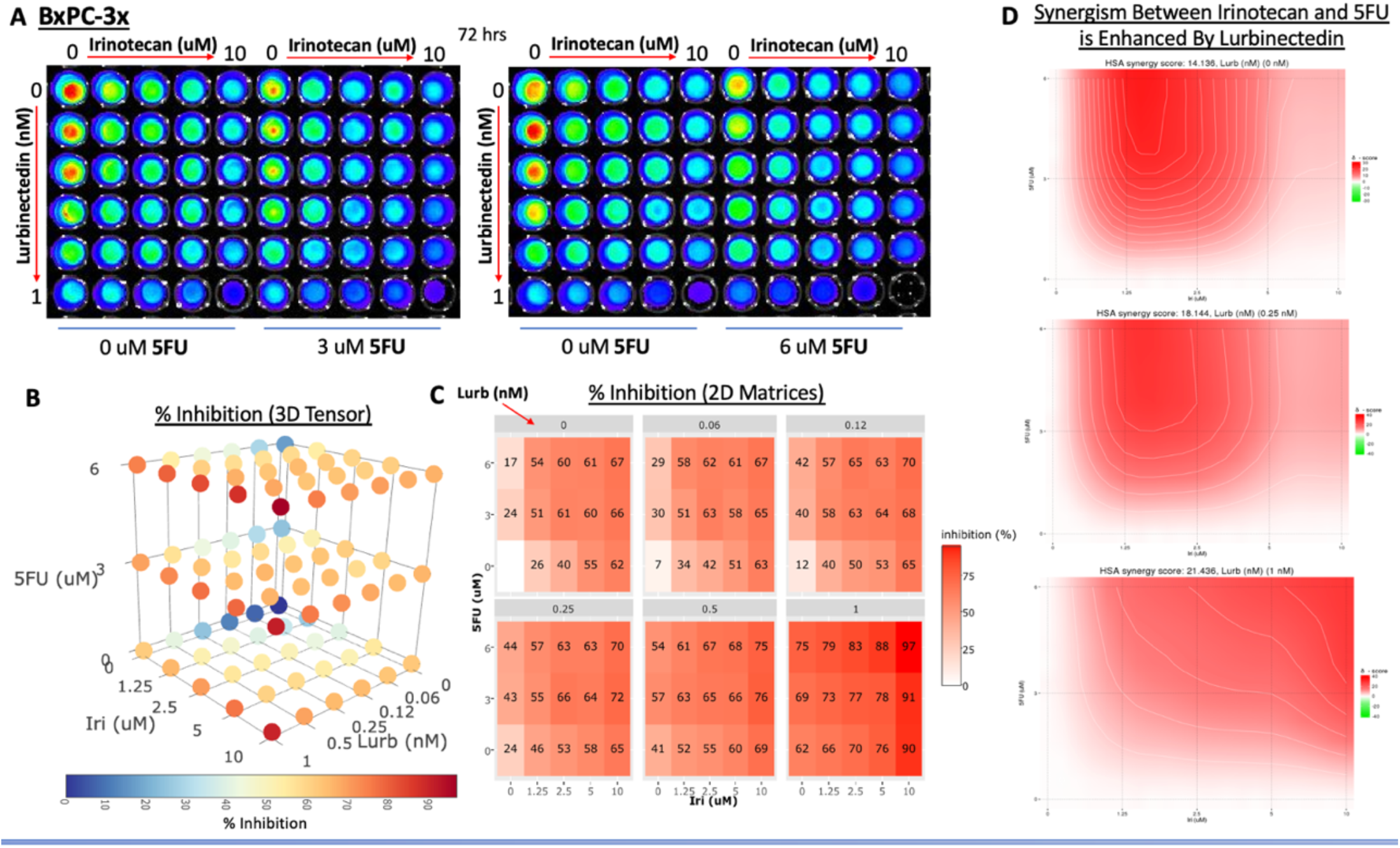
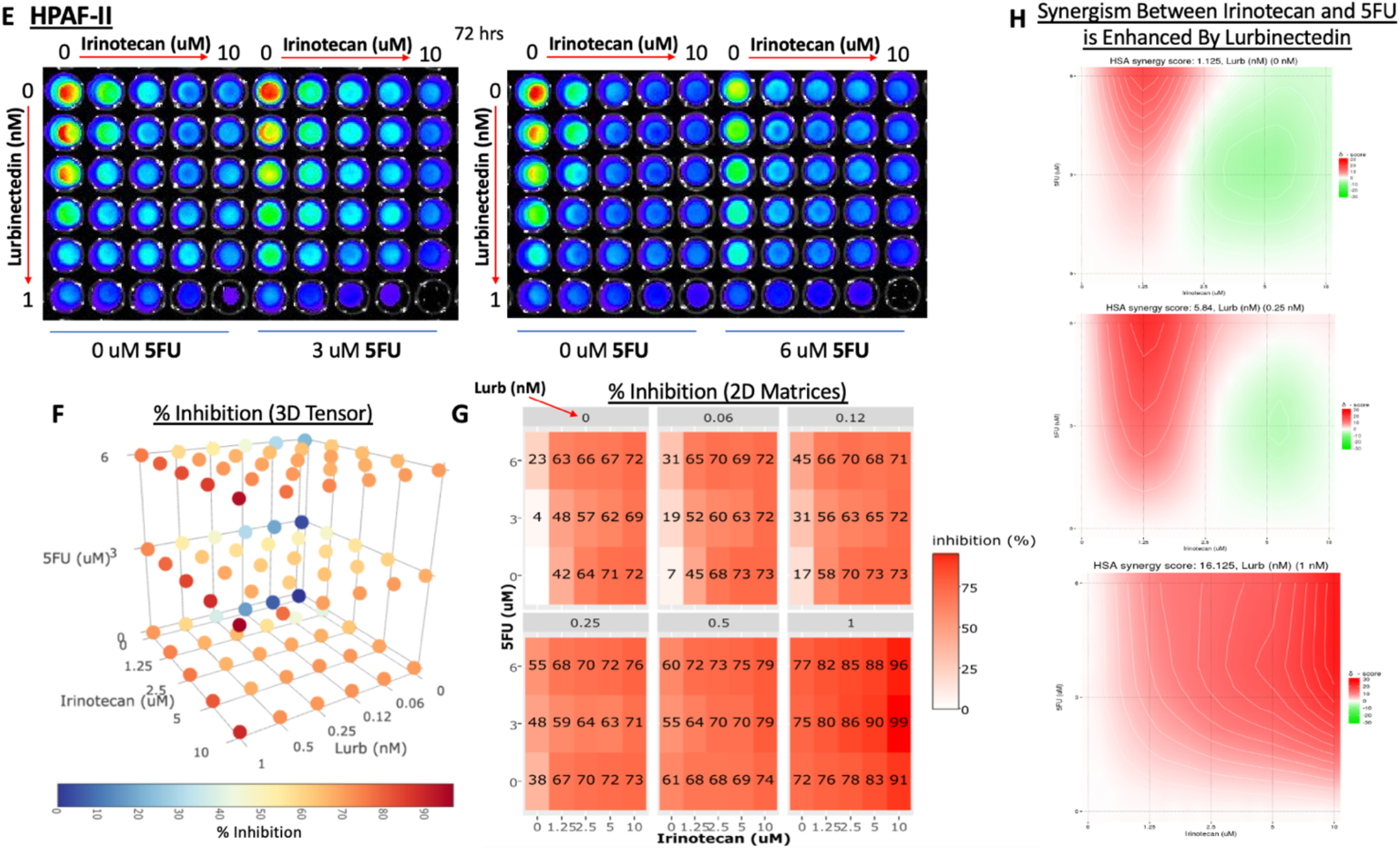
Triple combination cell viability assays of lurbinectedin, irinotecan, and 5-FU indicate that the three drugs act synergistically in killing pancreatic tumor cells. (**a**) BxPC-3 and (**e**) HPAF-II cell lines were treated with a triple combination of lurbinectedin, irinotecan, and 5-FU at varying concentrations for 72 hours. Following treatment, cells were visualized with CellTiterGlo technology, and synergism analysis was performed using SynergyFinder. (**b**) & (**f**) Percent inhibition was calculated and can be visualized in three dimensions on the 3D tensor plots, where darker colors correspond to higher percent inhibition. There is a clear trend across both cell lines that as each individual drug concentration increased, percent inhibition also increased. (**c**) & (**f**) Percent inhibition can be visualized in 2D matrices, where lurbinectedin is held to a constant concentration per matrix. These matrices confirm that the cell viability decreases as each individual drug rises in concentration. (**d**) & (**h**) Heatmaps display the HSA synergy scores between irinotecan and 5-FU at different concentrations of lurbinectedin. The average synergy score, corresponding to the average synergistic response due to the two-drug combination between irinotecan and 5-FU, is displayed above the heatmaps, with lurbinectedin concentration constant within each heat map yet differing across heatmaps. Without lurbinectedin, irinotecan and 5-FU had an average synergy score of 14.136 and 1.125 in BxPC-3 and HPAF-II, respectively. When 1 nM of lurbinectedin was added, the average synergy score between irinotecan and 5-FU increased to 21.436 and 16.125 in BxPC-3 and HPAF-II, respectively, demonstrating lurbinectedin is enhancing the synergism between irinotecan and 5-FU.

Overall, our results indicate that lurbinectedin is an efficient killer of human derived pancreatic cancer cell lines as a single agent and in combination with irinotecan and 5-FU, as the drug induces a variety of changes in protein and cytokine expression as a single agent and in combination with other anticancer drugs. Cytokine profiling and western blot analysis help reveal the mechanism behind lurbinectedin and irinotecan synergism, which is likely rooted in DNA damage, replicative stress, upregulation of death receptor ligands such as FAS, and a downregulation of Bcl-2.

## Discussion

Given the poor prognosis of advanced pancreatic cancer, it is imperative that better treatments are researched and eventually implemented in a clinical setting. In treating patient-derived pancreatic cancer cell lines with lurbinectedin as a single agent, lurbinectedin showed highly potent killing of such cells, with 72-hour IC-50s corresponding to sub-nanomolar concentrations in each cell line (**Figure 2A**). Irinotecan and 5-FU, which are commonly used in modern pancreatic cancer chemotherapeutic regimens, exhibited 72-hour IC-50s in the low micromolar range (**Figure 2B & Figure 2C**). Thus, lurbinectedin is much more potent by comparison.

After establishing lurbinectedin as a potential single agent killer of pancreatic tumor cells, lurbinectedin was combined with irinotecan to determine if the two drugs acted synergistically in killing pancreatic tumor cells. Such a hypothesis was put forward for a variety of reasons. Currently, lurbinectedin and irinotecan are being combined in a variety of tumor types in clinical trials. Additionally, lurbinectedin and irinotecan both act on DNA and are shown to induce DNA damage and breaks. With one drug that inhibits DNA repair and replication, and another that inhibits transcription, it was hypothesized to be a potent combination in killing pancreatic tumor cells, which heavily rely on replication and transcription relative to normal cells. Further, given that irinotecan is a rather standard drug in many chemotherapeutic regimens to treat pancreatic adenocarcinoma, and lurbinectedin is being tested as a monotherapy in a variety of advanced solid tumors (including pancreatic adenocarcinoma), there is potential for such a combination to have clinical relevance.

Our hypothesis was confirmed via two drug CTG cell viability assays and subsequent SynergyFinder analysis (**Figure 3**). Across all three cell lines, percent inhibition increased as the two individual drug concentrations increased, with the strongest inhibition corresponding to the bottom-right most corner wells of the 60-wells displayed having purple or black fluorescence. Such low fluorescence indicates little to no ATP detection following cell lysis, indicating these cells were no longer metabolically active and were likely dead. As indicated by the HSA Synergy Score matrices, lurbinectedin and irinotecan had synergy scores exceeding 15 in every cell line. The manufacturer of SynergyFinder indicates that synergy scores greater than 10 denote synergistic, as opposed to additive, killing of cells. As a byproduct of this synergism, the combination therapy of lurbinectedin and irinotecan allow for reduced concentrations of each drug individually while still maintaining a robust apoptotic response. Using lower concentrations of each drug individually could potentially reduce overall toxicity to animal models or patients, but more research is needed to confirm such a hypothesis.

Given the strength of the synergism between lurbinectedin and irinotecan, western blot analysis and cytokine profiling analysis were both performed to better understand the mechanisms underlying the synergistic killing of the cancer cells under the individual and combination drug conditions. As hypothesized, combination conditions across multiple cell lines showed an upregulation of γH2AX (**Figure 4B & Figure 4D**), a marker for DNA damage [24], indicating that genotoxicity and DNA breaks likely play a crucial role in the synergistic interactions between the two drugs.

Normally, the ATR/Chk1 pathway plays a role in protecting the genome from DNA damage and replication stress [25], and the pathway is activated as a cellular response to DNA damage [26]. Given that lurbinectedin and irinotecan’s mechanisms rely on inducing DNA damage and some replication stress, it was hypothesized that ATR and Chk1 expression would likely be upregulated by the drug combination. **Figure 4D** demonstrates that under combination therapy, PANC-1 demonstrates a relatively strong upregulation of Chk1 and a slight upregulation of ATR. Further, given that ATR is also seemingly slightly upregulated in BxPC-3 under lurbinectedin and irinotecan treatment (**Figure 4B**), it is likely that the ATR/Chk1 pathway is being activated to some extent under combination conditions. This provides further evidence that DNA damage and potentially replicative stress contribute to the observed synergism.

Additionally, Bcl-2 is normally a pro-survival protein [27], and slight downregulation of Bcl-2 under combination conditions (**Figure 4A & Figure 4C**) likely indicates that Bcl-2 family proteins play a role in the apoptosis of the pancreatic tumor cells. Cleaved PARP and ATF4 expression (**Figure 4A & Figure 4C**) were rather inconsistent across cell lines, and thus the information on these proteins and pathways is currently inconclusive.

Cytokine profiling further revealed the mechanisms underlying synergism. Across the cytokine panel, there were a variety of upregulated and downregulated factors under different treatment conditions in BxPC-3 (**Figure 5A**) and HPAF-II (**Figure 5B**). The heatmaps denote the fold-change from the treatment conditions relative to the control (no drug) group. Noticeably, VEGF was downregulated in single drug and combination conditions across both cell lines. VEGF often plays an important role in promoting angiogenesis in healthy humans [28], and lurbinectedin has been shown in previously to decrease VEGF [29].

Fas, a ligand involved in multiple apoptotic pathways [30], is upregulated across both cell lines under both monotherapy and combination conditions, which is consistent with expected results because lurbinectedin and irinotecan are efficiently inducing an apoptotic response in the cells. It is important to note that the effect of lurbinectedin on the tumor microenvironment is still under exploration and our results provide an incomplete yet novel understanding of the mechanism underlying synergy.

After establishing synergism between irinotecan and lurbinectedin and better understanding the mechanisms behind their synergism, it was hypothesized that a triple combination of lurbinectedin, irinotecan, and 5-FU might be a potent combination therapy capable of killing pancreatic tumor cells at relatively low concentrations of each individual drug. Given irinotecan and 5-FU are commonly used together, and lurbinectedin and irinotecan are being explored as a two-drug combination in clinical trials in many tumor types, it was important to test in the preclinical setting if adding lurbinectedin to a regimen containing irinotecan and 5-FU would be feasible. If such a drug combination can be affirmed in the preclinical setting, there could potentially be clinical application for such a triple combination.

Given the well documented antitumor activity of the three chemotherapeutics, it is rather unsurprising that **Figure 6** confirms that as each drug increases in dosage, higher percent inhibition is seen (indicating lower viability). Further, **Figure 6D** and **Figure 6H** demonstrate that the average synergism between irinotecan and 5-FU is enhanced by the presence of lurbinectedin. The presence of lurbinectedin is enhancing the efficiency by which 5-FU and irinotecan are capable of synergistically killing the cells, while lurbinectedin itself is interacting with the drugs in a synergistic fashion and contributing to lower cell viability. Our results suggest the possibility that lurbinectedin could have clinical relevance in a triple combination therapy with irinotecan and 5-FU.

Overall, the data indicates that lurbinectedin monotherapy and combination therapy with irinotecan or irinotecan and 5-FU are potent killers of BxPC-3, PANC-1, and HPAF-II. The two-drug combination seems to rely on DNA damage and replicative stress and an upregulation of cytokines related to apoptotic pathways. The triple combination therapy of lurbinectedin, irinotecan, and 5-FU demonstrates highly efficient and synergistic killing of pancreatic cancer cells, as lurbinectedin enhances the synergism between 5-FU and irinotecan. Our results are investigating the molecular mechanistic basis underlying the novel synergistic combination in pancreatic adenocarcinoma cells.

Future experiments will include mouse models in a tumor growth and mouse survival study to test whether lurbinectedin, irinotecan, and 5-FU monotherapy or combination treatments can reduce tumor size and prolonging lifespan. To better understand the mechanism behind lurbinectedin and irinotecan’s synergism, more biomarkers for DNA damage, replicative stress, caspase activation, and apoptosis need to be probed for via western blot analysis. Future experiments will also include co-culturing cytotoxic CD8+ T-lymphocytes as well as Natural Killer (NK)-cells with pancreatic tumor cells under monotherapy and combination therapy conditions to analyze whether the drugs are capable of sensitizing tumor cells to immune killing. Such experiments will provide a more robust analysis of the drug combination and shed further insights into whether the drug combination can modulate the tumor microenvironment. Such work will further support the translation of the combination in a clinical setting.

## Conclusions

Overall, the data indicates that lurbinectedin monotherapy and combination therapy with irinotecan or irinotecan and 5-FU are potent killers of human-derived pancreatic adenocarcinoma cell lines. The two-drug combination appears to rely on DNA damage and replicative stress and an upregulation of cytokines related to apoptotic pathways. The triple-combination therapy of lurbinectedin, irinotecan, and 5-FU demonstrates highly efficient and synergistic killing of pancreatic cancer cells, as lurbinectedin enhances the synergism between 5-FU and irinotecan. We expect in the future this therapeutic approach may be of use for patients with locally advanced or metastatic pancreatic cancer. Our results are developing a novel preclinical synergistic combination in pancreatic cancer with a feasible path to testing such a combination in a clinical setting.

## Author Contributions

Conceptualization, all authors; methodology, Tej Tummala, Andrew George, Ashley Sanchez Sevilla Uruchurtu, Kelsey E. Huntington, Lanlan Zhou, and Wafik S. El-Deiry; software, Tej Tummala and Wafik S. El-Deiry; validation, Tej Tummala, Lanlan Zhou, and Wafik S. El-Deiry; formal analysis, Tej Tummala and Wafik S. El-Deiry; investigation, Tej Tummala and Wafik S. El-Deiry; resources, Tej Tummala, Abbas E. Abbas, Christopher G. Azzoli, and Wafik S. El-Deiry.; data curation, Tej Tummala and Wafik S. El-Deiry; writing— original draft preparation, Tej Tummala; writing—review and editing, Tej Tummala and Wafik S. El-Deiry; visualization, Tej Tummala and Wafik S. El-Deiry; supervision, Wafik S. El-Deiry; project administration, Wafik S. El-Deiry; funding acquisition, Wafik S. El-Deiry. All authors have read and agreed to the published version of the manuscript

## Acknowledgments

W.S.E-D. is supported by the Mencoff Family Professorship at Brown University. W.S.E-D. is an American Cancer Society Research Professor.

## Appendix A

Some lanes were cropped out of Figure 4 due to irrelevance in this paper. The full Western Blot can be seen below. It is essential to note that no lanes were edited or altered to unfairly manipulate the data.

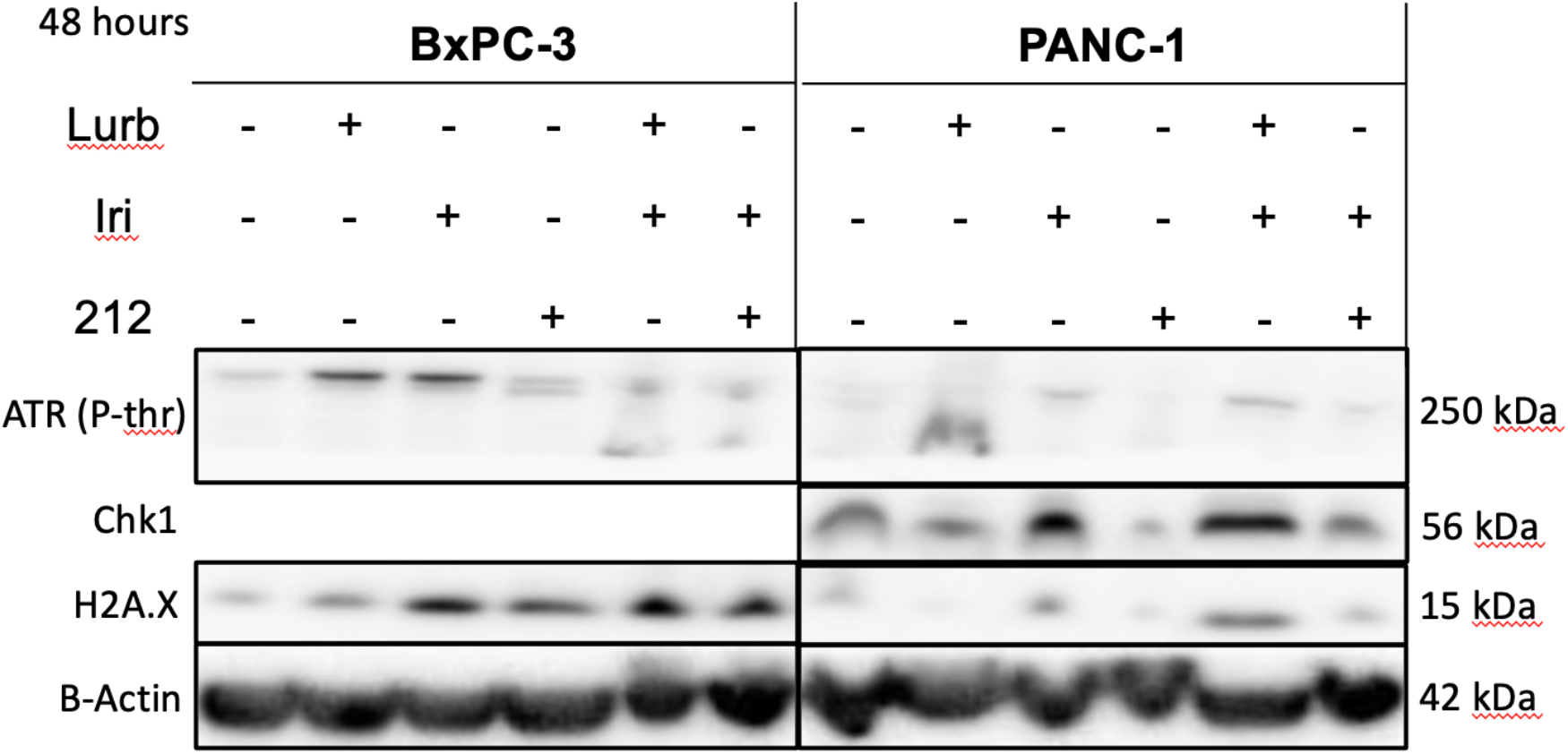

## Disclaimer/Publisher’s Note

The statements, opinions and data contained in all publications are solely those of the individual author(s) and contributor(s) and not of MDPI and/or the editor(s). MDPI and/or the editor(s) disclaim responsibility for any injury to people or property resulting from any ideas, methods, instructions or products referred to in the content.

